# PREMONition: An algorithm for predicting the circadian clock-regulated molecular network

**DOI:** 10.1101/463190

**Authors:** Jasper Bosman, Zheng Eelderink-Chen, Emma Laing, Martha Merrow

## Abstract

A transcriptional feedback loop is central to clock function in animals, plants and fungi. The clock genes involved in its regulation are specific to - and highly conserved within - the kingdoms of life. However, other shared clock mechanisms, such as phosphorylation, are mediated by proteins found broadly among living organisms, performing functions in many cellular sub-systems. Use of homology to directly infer involvement/association with the clock mechanism in new, developing model systems, is therefore of limited use. Here we describe the approach PREMONition, <u>PRE</u>dicting <u>M</u>olecular <u>N</u>etworks, that uses functional relationships to predict molecular circadian clock associations. PREMONition is based on the incorporation of proteins encoded by known clock genes (when available), rhythmically expressed clock-controlled genes and non-rhythmically expressed but interacting genes into a cohesive network. After tuning PREMONition on the networks derived for human, fly and fungal circadian clocks, we deployed the approach to predict a molecular clock network for *Saccharomyces cerevisiae*, for which there are no readily-identifiable clock gene homologs. The predicted network was validated using gene expression data and a growth assay for sensitivity to light, a zeitgeber of circadian clocks of most organisms. PREMONition may be used to identify candidate clock-regulated processes and thus candidate clock genes in other organisms.

## Introduction

The circadian clock is a ubiquitous biological program^1^. It creates a temporal structure that anticipates the regular and predictable changes in the environment that occur each day^2^. Circadian clocks are cell-based and are found in organisms from all phyla. The molecular leitmotif of the eukaryotic circadian clock involves transcription factors that activate a set of genes that encode proteins capable of down-regulating their own expression, forming a negative feedback loop. However, recent experiments suggest that circadian clock properties persist in the absence of identified clock genes. For instance, *Neurospora crassa* shows free-running rhythms and circadian entrainment without the clock gene *frequency*^3,4^ and rhythms in the redox state of peroxiredoxin persist in many model genetic systems in the absence of transcription or of clock genes^5^. Paracrine signaling can sustain circadian rhythms in pacemaker (neuronal) tissue that lacks clock genes and displays no endogenous rhythm^6^. One interpretation of this is that feedback integral to circadian clocks spans multiple molecular ‘levels’, from transcription to metabolism. We conjecture that this multi-level response can be captured through functional relationships, referred to as functional interactions. We know of no models that explicitly attempt to integrate these different levels to more fully describe the mechanisms of the circadian clock. Here, we describe the method PREMONition, PREdicting MOlecular Networks, to construct multi-level (functional) molecular clock networks using publicly available datasets and bioinformatics tools. After tuning the method on clock genetic model systems, we deployed PREMONition to predict a molecular clock network in *S. cerevisiae*, an experimental system with a circadian phenotype that is not suited to genetic screens^7^. We validated the results of the yeast experiment *in silico* and *in vivo*. Methods like PREMONition, that span multiple levels in the cell, are essential to developing integrative models of complex biological processes.

## Results

### PREMONition applied to model circadian mechanisms

To construct circadian molecular networks PREMONition incorporates two sets of data: rhythmically expressed genes and evidence-based (physical and functional) interactions of molecules derived from published literature and stored in the STRING database^8^. The set of rhythmically expressed genes (clock-controlled genes (ccgs)) is derived from transcriptome analyses. Clock genes - those that have been determined to be involved in the central clock mechanism – are also included here, when known. By logic, clock genes must be rhythmically expressed at some level, whether it be RNA or protein expression or functional competence (e.g. determined by rhythmic post translational modification). Ccgs are genes that are controlled by the central clock mechanism that are not thought to feed back into the clock mechanism (no evidence that their disruption affects the clock). For the second data set, interacting proteins catalogued and/or predicted by STRING are increasing in number based on the massive amount of genome-wide quantitative measurements that are now common.

### Network construction

Within STRING molecular interaction networks are predicted and displayed. The network reflects the specified stringency of the statistical probability of interactions. One can thus make a network that is more or less connected and/or one that is larger or smaller by specifying statistical parameters. To develop the PREMONition method, we first created a molecular interaction network formed of proteins encoded by canonical clock genes only. As expected, these proteins, known to be involved in circadian transcriptional feedback loops in human cells, *D. melanogaster* and *N. crassa*, form small, highly connected networks (Figs. 1A, B and C). However, when combining clock genes together with ccgs derived from published transcriptome analyses^9–11^ (Table S1), the derived networks comprised disjointed sub-networks rather than a cohesive network (Figs. 1D, E and F). For the human network, only two interactions between the clock genes and ccgs were identified.

**Figure 1.**
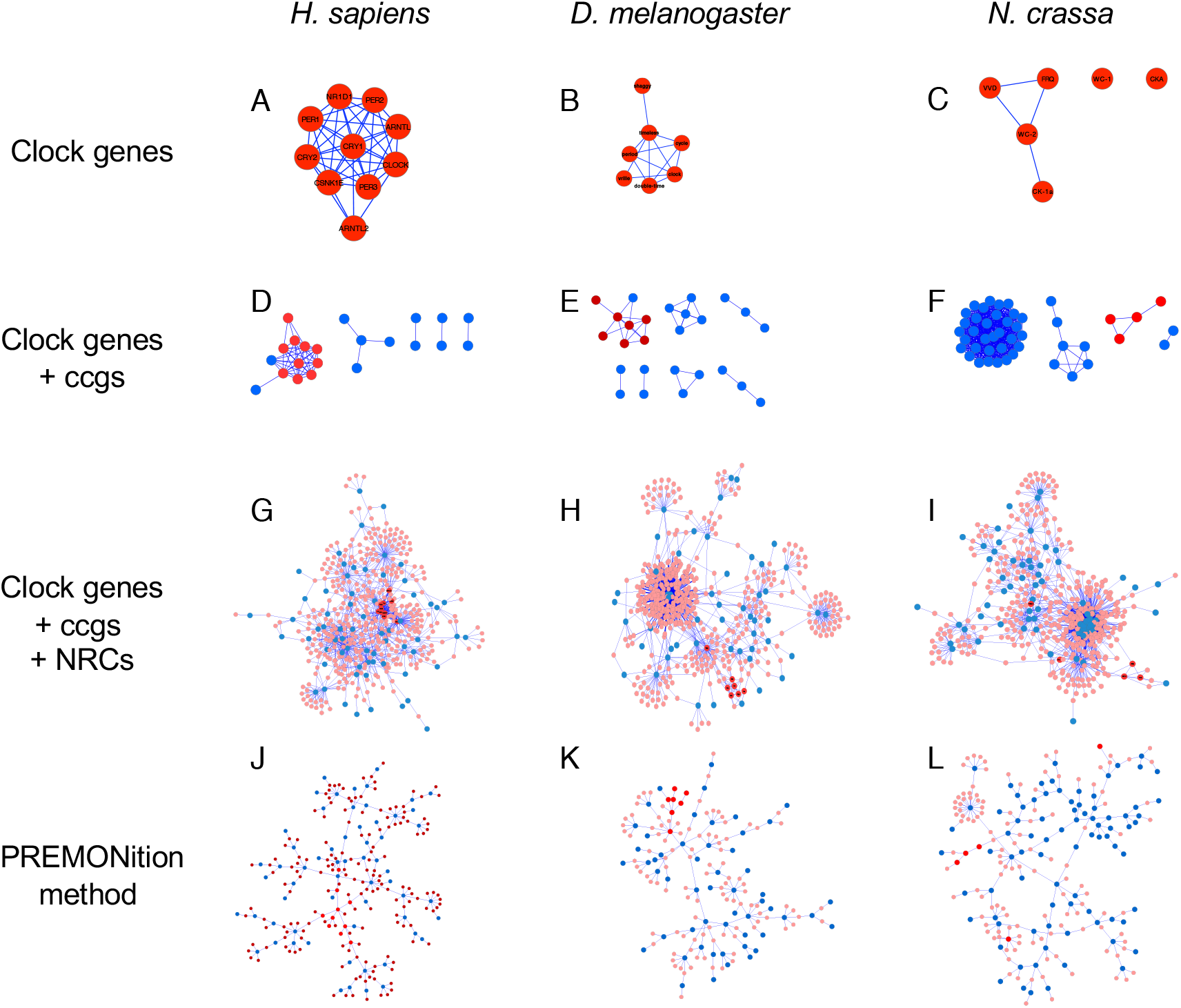
Molecular interaction networks comprising molecules encoded by human, *Drosophila* and *Neurospora* clock genes, ccgs and NRCs. Clock genes alone tend to form connections (A, B, C). Addition of ccgs, does not lead to a cohesive network (D, E, F). Addition of NRCs makes a highly connected network (G, H, I). Removal of the least probable NRCs connecting two rhythmic nodes and removal of terminal NRCs (panels J, K and L). Molecules that fail to be incorporated into the network in panels D-L are not shown. Spheres marked with red indicate proteins encoded by clock genes; blue spheres indicate proteins encoded by clock-controlled genes; pink spheres indicate proteins encoded by non-rhythmic connectors.

Forming a molecular interaction network associating all clock/clock-controlled genes is a key step in moving beyond studies that concentrate on single genes and their functions. It is well known that rhythmic RNAs can lead to non-rhythmic proteins and non-rhythmic RNAs can lead to rhythmic proteins^12–14^. Furthermore, post-translational modifications can confer rhythmic function even in the absence of rhythmic protein expression^14,15^. We therefore allowed PREMONition to incorporate <u>n</u>on-rhythmically expressed <u>c</u>onnectors (NRCs) into the ‘clock gene + ccg molecular interaction network. NRCs by definition are molecules that interact, physically or functionally, with molecules associated with clock genes and/or ccgs; as their name suggests, NRCs are not themselves identified as rhythmically expressed at the gene expression level. In the experiments described here, a relatively stringent probability score of ≥0.7 was used for *Drosophila* and human and ≥0.6 for *Neurospora* within STRING. This cutoff was derived by determining when highly interconnected networks would form, and when they would break (become fragmented) if higher scores were used. Networks were formed that incorporated 70%, 38% and 57% of ccgs from human, *Drosophila* and *Neurospora* gene sets, respectively (Figs. 1G, H and I). Networks were trimmed by allowing only a single NRC between two nodes (either clock gene or ccg; Table S2). The most probable connection was retained and all others were discarded. Through this process, which selects for the most likely interactions, networks were reduced from 483 to 248, from 498 to 227, and from 566 to 196 without decreasing ccg incorporation for the three species (human, *N.crassa* and *D.melanogaster* respectively). In all three organisms, an extensive and inclusive network was generated (Figs. 1 J, K and L).

### Network interpretation

The three PREMONition derived networks shared thirty-eight common Gene Ontology (GO) terms^16^. Ten of these terms were significantly enriched biological processes in the human, fly and *Neurospora* networks (Fig. S1 and Table S3). A thousand Monte Carlo simulations using random sets of genes, representing random networks the same size as the PREMONition derived networks, yielded not a single over-represented GO category. Applying GO analysis to each of the collections of ccgs also yielded no significantly enriched categories. Conversely, the GO categories that were detected in the analysis of entire PREMONition networks represent a large percentage of categories for a given genome. As such, these categories are not predictive – they cover too much of the genome to be useful for identification of discrete genes and proteins involved in clock function.

Orthology between the molecular networks was assessed using the InParanoid8 database^17^. For human, fly and fungus the 283, 267 and 245 unique molecules, referenced by their corresponding gene name were submitted to the InParanoid8 database, respectively, for a cross comparison. Table 1 shows the number of orthologous protein pairs, including orthologous clusters identified by Inparanoid 8 (Table S4). In total 30, 23 and 21 human, fly and *N. crassa* genes were found to be in common between the respective networks.

**Table 1.** Identification of orthologs with InParanoid8. The number of D.*melanogaster* (fly) orthologs that occur *H.sapiens* (human) and *N.crassa* (fungus) and between human and fungus is shown. Several of the orthologs are included in the PREMONition networks for the specific organisms. P-values indicate the significance of ortholog enrichment in the networks.

|  | **Orthologs** | **Networked** | **p-value** |
| --- | --- | --- | --- |
| **Fly - human** | 10711 | 19 | 2.08e-22 |
| **Fly – mold** | 4004 | 11 | 4.23e-28 |
| **Human - mold** | 5155 | 14 | 1.15e-33 |

### A yeast PREMONition network

Although clock properties such as systematic entrainment, indicative of a *circa* 24 h oscillator, can be elicited in yeast^7^, the clock phenotype is not robust enough to support a high-throughput genetic screen that could uncover candidate clock genes. There are no clock gene candidates in yeast that are predicted by homology searches or functional assays. In order to generate a putative circadian molecular interaction network for yeast, we used PREMONition. A yeast transcriptome was generated covering the first day following release into constant conditions (Fig 2A). One-hundred-and-thirty-seven genes were determined to be rhythmically expressed, with all phases of expression observed over the subjective day (Fig. 2B and Fig. 3). This set of ccgs did not form a fully interconnected network (probability score of >= 0.7) when submitted to STRING. Only 23 genes/proteins were connected. Allowing NRCs, however, resulted in a network of 477 interacting molecules, consisting of 123 rhythmic genes and 354 NRCs (Fig. 4). 14 rhythmic genes could not be integrated into the network. Referring to the networks described for human, *Drosophila* and *Neurospora* clock systems, 19, 11 and 19 genes, respectively, have orthologs in *S. cerevisiae* and 14 of those were identified in the yeast network.

**Figure 2.**
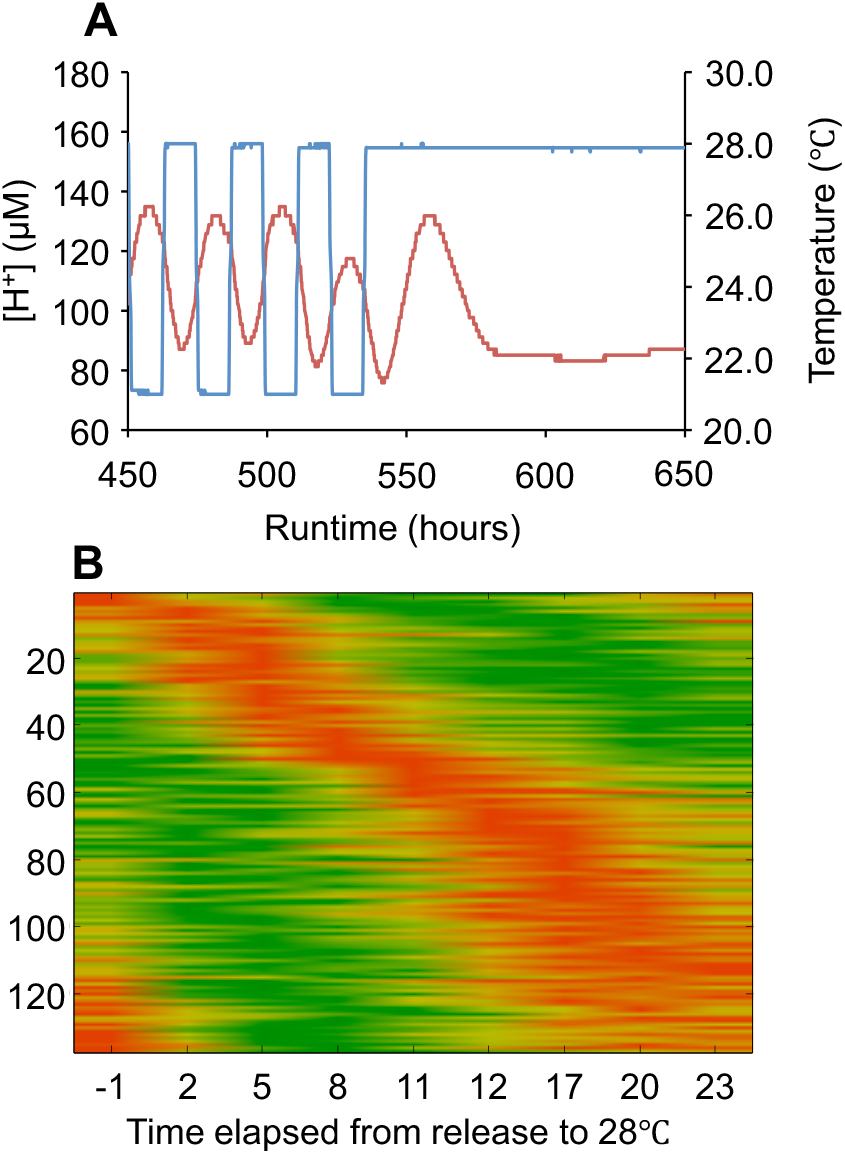
Rhythmicity of gene expression in yeast. Fermentor cultures were established^7^ using 24h zeitgeber cycles (21 °C / 28 °C, blue line) to synchronise the cells, as measured by pH of the culture medium (red line). Once stable synchronisation was observed, the temperature was held at 28 °C leading to one >24h oscillation in the pH (A). Cells were harvested every 3h for 24h, starting 1 hour before the last temperature transition (from 21 °C to 28 °C). The RNA transcriptome was analysed using microarrays, yielding a set of 137 rhythmic genes (p-value <0.01). The heat map shows the time of high expression in red and the time of low expression in green (B). Each individual signal is normalized between 0 (green) and 1 (red). They are plotted in order of peak phase.

**Figure 3.**
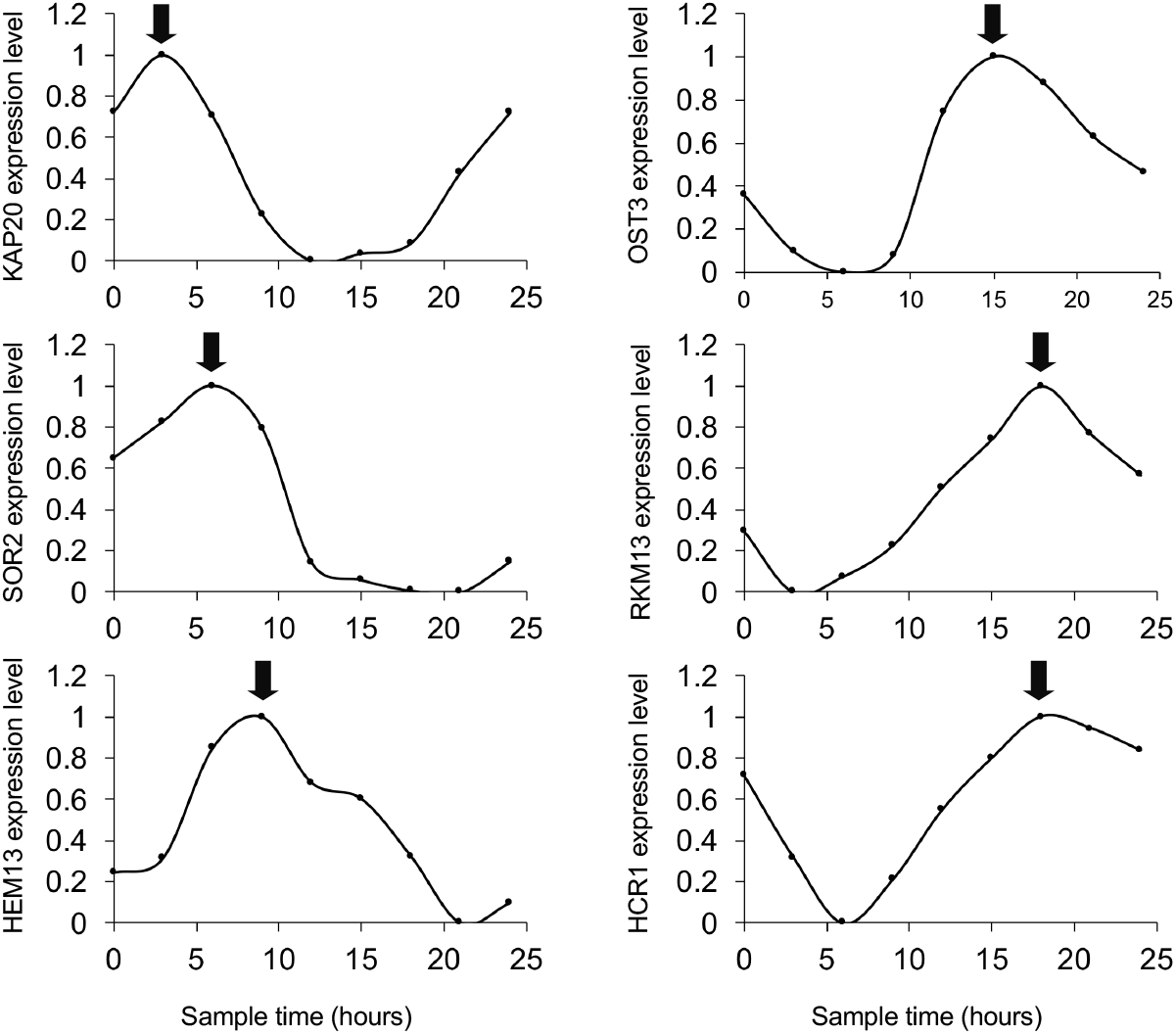
Rhythmic genes in yeast span ‘all’ circadian phases. The 137 rhythmic genes were grouped into 6 bins of 4 hours each based on their peak phase of gene expression. For each bin, the gene showing highest significance is shown. An arrow marks the peak phase of each gene. All time series were normalized such that their values fall between 0 and 1.

**Figure 4.**
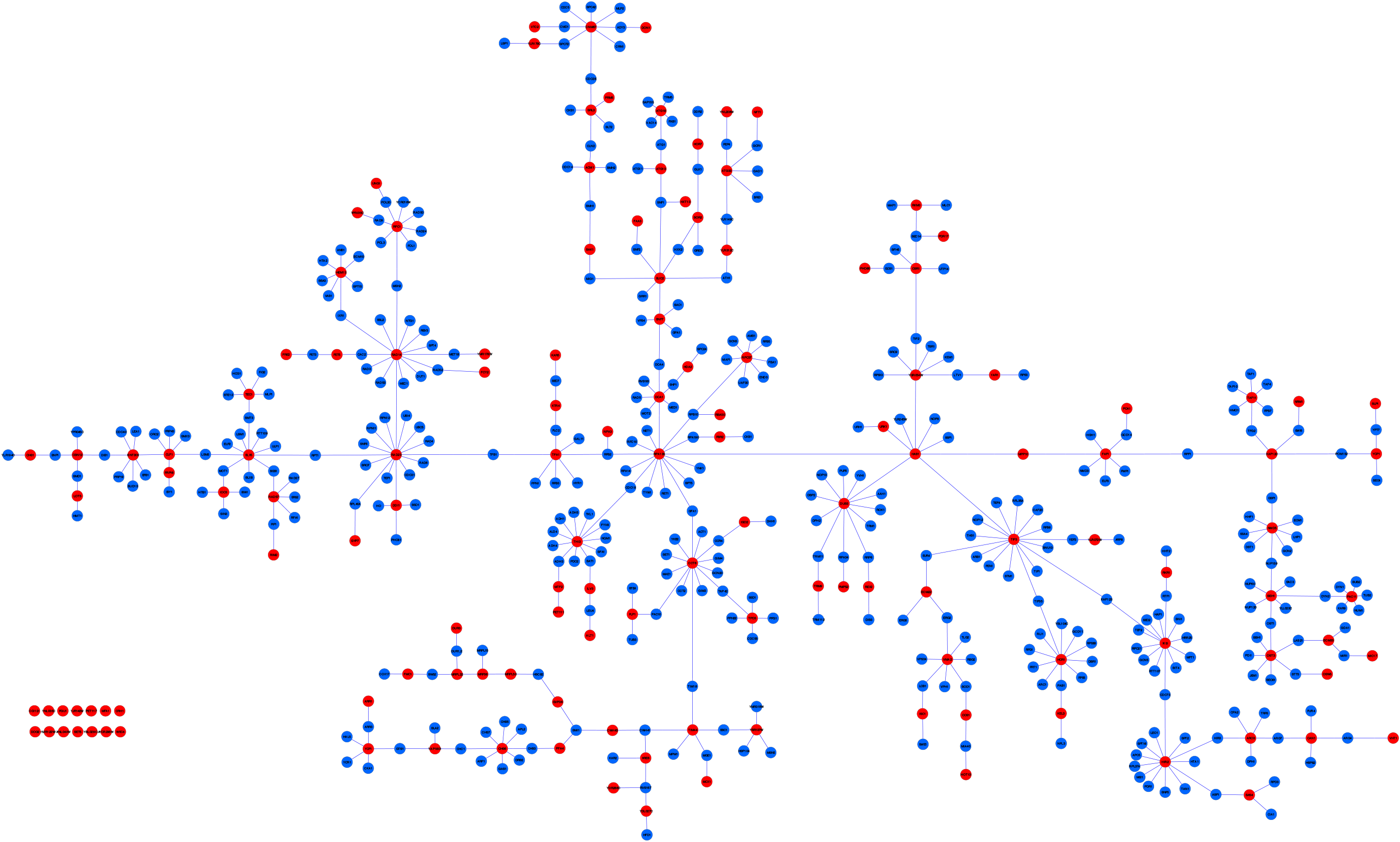
The PREMONition network for yeast. Proteins encoded by rhythmically expressed genes (red spheres) and their non-rhythmic connectors (blue spheres) are shown. The network is composed of 123 rhythmic genes and 354 NRCs, connected by 476 edges. 14 rhythmically expressed genes are disjoined from the network. The figure is scalable such that each gene/protein can be identified.

### Validation of the yeast PREMONition network

We attempted to experimentally validate the putative yeast circadian molecular interaction network in two ways. First, we relaxed the stringent threshold used to identify ccgs in our yeast transcriptome from a p-value of 0.01< to <0.05. This increased the number of ccgs from 136 to 526. We found that thirty-three of the 355 original *S. cerevisiae* NRCs were considered to be rhythmically expressed with the relaxed p-value (Fig. 4). This number (33 of 355) is higher than expected by chance (hypergeometric test p-value = 2.53^e-06^) and indicates that, as a method, PREMONition is able to identify clock associated molecules.

A second validation method was developed based on the observation that circadian clocks of surface-dwelling organisms are sensitive to light. Photobiology in yeast is unknown except for reports of suppression of growth in constant light conditions apparently via interaction with cytochromes^18,19^. We hypothesized that mutations in genes involved in circadian clock function will show either an increased or decreased sensitivity to light in a growth assay. We therefore assayed a selection of mutants carrying knockouts of genes from the network for their sensitivity to light. Five genes present in the PREMONition network were selected based on their relationship to known circadian systems across different phyla. SOD1 (YJR104C) encodes a cytosolic copper-zinc superoxide dismutase, having a rhythmic mRNA expression level in mouse liver^20^. Superoxide dismutase activity is rhythmic in the dinoflagellate *Gonyaulax polyedra*^21^ and the *sod–1* mutant in *N. crassa* shows changes in the period of circadian rhythms and abnormal responses to light as a zeitgeber for entrainment^22^. The yeast FET3 (YMR058W) protein is a multicopper oxidase, a family that is conserved in many organisms and showing circadian rhythms in *A.thaliana*^23^. Additionally, copper functions as a redox active cofactor for many proteins like cytochrome-c-oxidase and copper-zinc-superoxide dismutase (Cu/Zn-SOD)^24^. Its expression is rhythmic and is important for circadian rhythms in the mammalian pacemaker^25^. The SGS1 (YMR190C) protein is a RecQ-family nucleolar DNA helicase. RecQ mRNA is among the few transcripts that cycle independent of Kai clock proteins in cyanobacteria (*S. elongatus*)^26^. The CHC1 (YGL206C) gene encodes a Clathrin heavy chain protein broadly involved in intracellular protein transport and endocytosis. Clathrin-interacting proteins have been implicated as potential clock effectors or as clock regulated genes and proteins in flies and in mice^27,28^. HOG1 (YLR113W) is the homolog of mitogen activated protein kinase (MAPK). Many MAPK proteins are involved in circadian regulation in the mammalian clock^29^. In *Neurospora* it functions as a mediator of clock-regulated outputs^30^. HCR1 (YLR192C) is an elongation initiation factor regulating translation. This gene scored amongst the most highly rhythmic (p = 0.0001) and its protein performs an important cellular function that has relatively rarely been investigated for clock function^31^. We used knockouts in all of these genes in an assay for suppression of growth by light (Fig. 5). With the exception of HCR1, all knockouts tested showed a significant alteration in growth suppression by light. One of these, a DNA helicase homolog, was resistant the effects of light. The other strains – knockouts for genes encoding SOD1, HOG1, CHC1 and FET3 – all showed more growth suppression in the presence of light than did the parental yeast strain.

**Figure 5.**
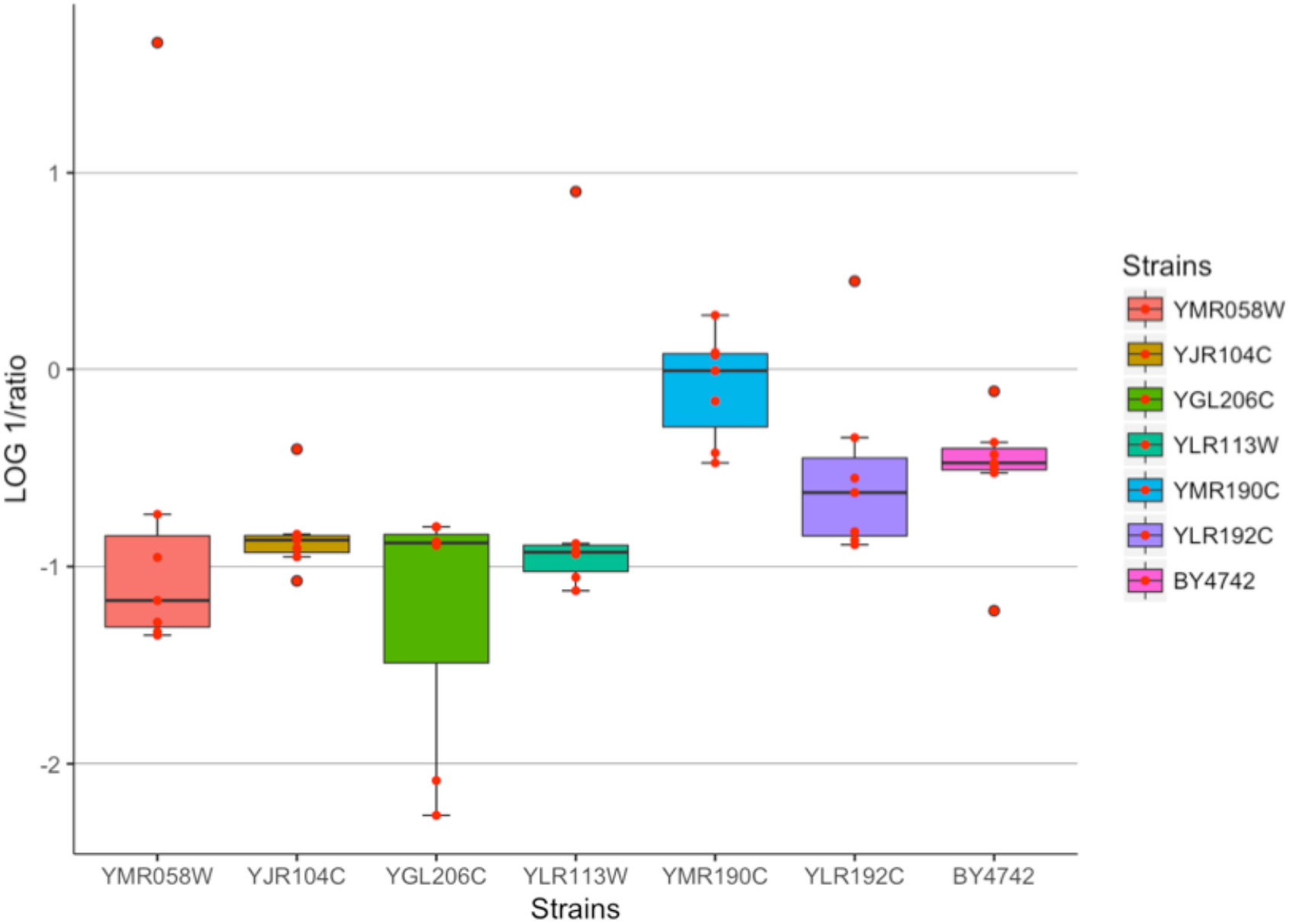
Light mediated growth repression of yeast. Growth in darkness was compared to growth in constant blue light at 15 °C. Single experiments are indicated by red spheres. All strains except YLR192C are significantly different relative to the control.

## Discussion

The circadian clock is often referred to as a success story for the field of behavioural genetics. Genetic screens and molecular genetic technologies provided breakthroughs in identification of clock genes^32–35^, loosely defined as genes without which free running circadian rhythms are disturbed or absent. The validated clock genes themselves encode highly networked transcription factors and kinases. In addition, collections of ccgs for many organisms and tissues have accumulated in public databases and can be used in diverse ways to generate hypotheses concerning clock regulation and clock regulated physiology^36^. The ccgs, by definition regulated directly or indirectly by the circadian clock mechanism, have rarely been directly linked to said mechanism. In addition, substantial experimental evidence has accumulated concerning non-transcriptional levels of circadian clock regulation yet this concept is not yet integrated into many models of clock mechanisms. Indeed, puzzling evidence of non-transcription-based clocks have been published. One interpretation of these observations is that embedding circadian clocks at diverse levels will enable their function under diverse conditions, thus fulfilling the prediction that the circadian clock should be adaptive.

How to model this? Combining and integrating information from different levels can be done by using machine learning techniques. The putative clock protein CHRONO was identified by combining time course results, interactions, ubiquitous expression and sequence homology using Naïve Bayes learning^37^. Indeed, machine learning typically depends on well characterized datasets to train the algorithm, e.g. to predict protein interactions using text mining with support vector machines^38^. Based on the data resources used and contextual understanding of the algorithm, machine learning could also assign cross-level relationships. However, different data resources require different approaches in data processing because using a single data resource may bias results.

Some data repositories collect data from ‘all’ functional and/or association interactions, including proteins, genes, transcripts, metabolites, etc. and, as these resources mature, they can be mined for insights into how a given network functions in a given tissue. We thus developed an algorithm called PREMONition that uses a set of known clock genes and ccgs in conjunction with various resources encapsulated within the STRING database (e.g. SMART, KEGG, etc.) to construct a plausible circadian molecular interaction network that can be used as a springboard for hypothesis generation and validation.

### The PREMONition tool

There are several algorithms that have been used to predict networks. Breadth First Search (BFS)^39^ visits all child nodes before making extension nodes. Depth First Search (DFS)^40^ first goes into depth of a single child before going on the next child. Depth Limited Search (DLS) uses DFS with a limited search space and a fixed search depth^41^. It increases the search depth until a solution is found, within the search space. Iterative Deepening Depth First Search (IDDFS)^41^ is similar to DFS, but continually expands its search depth until a solution is found. PREMONition uses a DFS implementation, since it makes an uninformed (i.e. all edges have the same probability), fully interconnected network. However, as in DLS, it uses a limited search space (a fixed number of NRCs are allowed between the ccgs). When the search limit is reached, the algorithm reverts to its parental node and deepens other child nodes.

PREMONition exploits the power of curated, precomputed functional interaction data (e.g. protein-protein interaction inferred from screens or shown in targeted experiments). It is an approach, with similarities to Biana^42^, POINeT^43^, SNOW^44^ and UnuHI^45^, that can be used to construct a network. With the publication of genome-wide, protein interaction databases, several methods to predict the proteome-wide interactomes have been published^46,47^. PREMONition uses many of the same resources as other methods but aims not to predict an interactome that incorporates the entire proteome, on one extreme, nor one that involves only a single input protein, on the other. Rather, we wish to form an interaction network around the molecules encoded by clock and clock-regulated genes. We achieve this by allowing the insertion of single NRCs (linker molecules). Increasing the number of NRCs, similar to expanding the search limit in DLS, will also increase the “noise” (non-specific or less specific interactions that would thus be more likely to be incorrectly assigned) in the network. In contrast to BFS, DFS, DLS and IDDFS, PREMONition tries to decrease noise and to increase the representation of more likely network components by removal of all but the most statistically significant interactors via an iterative process of sequential edge removal of the least probable interactor. On one hand, this makes visualization and analysis of the interaction network simpler. On the other hand, NRCs and their pruning also effect network parameters such as connectivity, degree distribution, clustering coefficient, characteristic path length network diameter and between-ness. It is striking that the set of rhythmically expressed components – ccgs with their accompanying experimental evidence - fails to form a cohesive network unless NRCs, based on their own experimental evidence, are integrated. The resulting networks derived for different organisms are functionally similar based on GO analysis (Fig. S1 and Table S3) despite the fact that input datasets are distinct.

The PREMONition algorithm is adaptable. Of the two input components, both can be modified for stringency. The set of rhythmic genes can be increased or decreased according to statistical thresholds. It could also be limited to those that oscillate with a certain phase to predict time-of-day-specific molecular networks. Or based on a set of cell/tissue/organ specific expressed genes to predict clock-regulated processes that are relevant for the distinct function of cells/tissues/organs^48^. Similarly, the NRCs can be increased or decreased in number by changing statistical threshold of an interaction or by altering the allowed number of connections. Alternative or additional datasets could be used as source data, including those for non-circadian biology, to predict non-clock molecular mechanisms. (Circadian clocks modulate expression over time of day but in clock-less mutants these genes are typically also expressed.)

A limitation of the PREMONition method is that it requires a mature interaction database (we used STRING exclusively for these experiments). The formation of the network thus depends on the quantity and quality of the annotation of the interactions. In emerging model systems or those that are less studied, there will simply be less data available thus leading to a sparse or less accurate network. With respect to the models that we used, the STRING database contains 3,364 *N.crassa* proteins connected with 24,332 interactions with a probability score ≥ 0.70 and an average confidence score of 0.84 (on average 7 connections per molecule). *D.melanogaster* is represented with 5,598 proteins connected with 93,915 interactions and an average confidence score of 0.89 (on average 16 connections per molecule). This roughly reflects the 7 times more publications in PubMed for *D. melanogaster* relative to those for *N. crassa.* Further, the network composition will be biased toward the most successful research for that system: if an organism has been primarily used to study enzymatic pathways, then the archived data – and the network formed from it - will be fundamentally different than that for an organism that has been used to study development. A disadvantage of using the STRING database is that it merely provides a probability of interaction in a static network, whilst the true interaction probability may be time-of-day dependent^49^, as well as condition or tissue dependent.

### Insights into the clock in yeast based on PREMONition network topology

We identified the most densely connected components of the reconstructed yeast network, reasoning that they may suggest organism-specific regulatory mechanisms (Table S5). Several of these densely-connected proteins are involved in (generic) transcription and translation processes (TIF3, the translation initiation factor eIF-4B; HCR1, part of the eIF-3 complex; RPA135, the RNA polymerase; RPC37, a subunit of RNA polymerase complex; TFA1, a recruiter of polymerase II; IKI3, a subunit of the elongator complex; and HAS1, an ATP dependent RNA helicase). These, *per se,* are not likely first-line clock components. HIR2 also regulates transcription, as part of a complex regulating nucleosome assembly. This protein recruits SWI-SNF, which plays an important role in nucleosome remodeling and carbon source switching^50^. SNF1, together with HOG1, whose homolog is clock regulated in *N. crassa*^51^, and BMH2, regulates MSN2/4, which in turn regulates the expression of PRX1^52^. The PREMONition yeast network thus maps out a possible mode of how the clock regulates PRX, whose rhythmicity in oxidation state is observed in eukaryotes, prokaryotes and even in the archeae^53,54^.

### Insights based on comparing networks

We compared the genes which are common in the human, *D. melonogaster, N.crassa* and yeast networks (using GO and Inparanoid) revealing 13 genes common to all four networks (Table 2). Of these, two genes are rhythmic in yeast with a p-value < 0.05 (YDR188W (CCT6) and YPR173C (VPS4)). The remaining 12 would not have been included in this dataset without incorporating NRCs. Clustering all 13 genes using PANTHER^55^ provides insights into their functions. Ten of these genes are involved in nitrogen compound transport: *CDC5, ACT1, RPD3, RPN11, HRR25, HOG1, EFB1, CDC19* and *TKL1*. We observed clock regulation of an ammonium and an amino acid permease in yeast under conditions permissive for circadian entrainment^7^ and nitrogen metabolism and utilization is known to be regulated by the circadian system in prokaryotes (the cyanobacterium *Synechococcus* fixes nitrogen during the night^56^) and in eukaryotes at multiple levels in many organisms (e.g.^57^). Three genes shared by all of our networks are involved in vacuole organization: *YPT7, ACT1* and *HRR25*. Of these, *HRR25* encodes a Casein kinase I homolog with many diverse functions in most organisms. Casein kinase plays a major role in the mammalian circadian clock through phosphorylation of the PERIOD proteins and BMAL1^58^ and core clock proteins in *D.melanogaster*^59^.

**Table 2.** Yeast genes included in the yeast PREMONition network which have overlapping gene ontology terms and orthologs in the human, fly and fungus networks.

| **Systematic name** | **Standard name** | **Name description** |
| --- | --- | --- |
| YNL330C | RPD3 | Reduced Potassium Dependency |
| YPL204W | HRR25 | HO and Radiation Repair |
| YDR188W | CCT6 | Chaperonin Containing TCP-1 |
| YIL021W | RPB3 | RNA Polymerase B |
| YML001W | YPT7 | Yeast Protein Two |
| YFL039C | ACT1 | ACTin |
| YAL003W | EFB1 | Elongation Factor Beta |
| YLR113W | HOG1 | High Osmolarity Glycerol response |
| YMR001C | CDC5 | Cell Division Cycle |
| YPR173C | VPS4 | Vacuolar Protein Sorting |
| YPR074C | TKL1 | TransKetoLase |
| YAL038W | CDC19 | Cell Division Cycle |
| YFR004W | RPN11 | Regulatory Particle Non-ATPase |

Four genes (*CDC5, HRR25, HOG1, CDC19*) encode proteins with kinase activity, a group of proteins known to be involved in the regulation and functioning of the circadian clock^60^. Yeast *HOG1* is an ortholog of *N.crassa cka* and hog-1, of which *cka* is known to be involved in its circadian clock^61^. When it is mutated Neurospora circadian rhythms are abolished^62^. *HOG1* is also a homolog of *OS-2* mitogen-activated protein kinase (MAPK). In *N.crassa* there is a feedback loop between the circadian clock and the MAPK pathway^30^. *N.crassa* encodes two other MAPK genes; MAK-2, a homolog of FUS3, and MAK-1, a homolog of SLT2 (YHR030C) and shaggy (*D. melanogaster*) which is a key player in the fly circadian clock through phosphorylation of TIMELESS^63^. *SLT2* is included in the yeast PREMONition network as an NRC (p-value for transcriptional rhythmicity = 0.74). *RPN11* is a homolog of Rpn11 (DmS13) in *D. melanogaster*. In *Drosophila*, light-dependent CRY degradation is regulated by the proteasome of which DmS13 is a subunit^64^. Genetic screens in mice have also yielded genes involved in the proteasome as key circadian clock regulators^65^. *CCT6* is homologous to the human *PSMD5*, a proteasome component, and acts as a chaperone during the assembly of the 26S proteasome. Interestingly, this protein shows interaction, via UBB and UBC, with CRY1. *CRY1* encodes a flavin adenine dinucleotide-binding protein, a key clock protein in mammals. *VPS4* is an ortholog of *VPS4B* in humans and *VPS4* (NCU06942) of *N.crassa*, both vacuolar protein sorting-associated proteins. Although no direct relationship between *VPS4* and the circadian clock have been reported, several suggestive physiologies are impacted by decreased function of VPS4. These include: increased heat sensitivity (possibly changing sensitivity to a zeitgeber), decreased replicative lifespan, chronological lifespan and decreased rate of respiratory growth (possibly related to the phenotype used to demonstrate circadian entrainment in yeast^66^)^67–69^.

The PREMONition NRCs shared by three networks thus point to conserved clock-relevant pathways. However, the number of candidate clock-involved genes regulating those pathways in yeast is drastically decreased by our algorithm.

### The yeast PREMONition network in general

Whilst a large proportion of the PREMONition network components in yeast do not have a direct homolog in the networks of the other three species, they do have homologous function. The RAD23 and RAD14 proteins bind damaged DNA, reminiscent of a documented clock function in mammals and the green yeast, *Chlamydomonas*^70,71^. CNM67, a hub with 16 connections in the yeast PREMONition network, is involved in spindle body formation, a connection to the cell cycle machinery, linking this network to observations of gating of cell division^72,73^. A subunit of the ubiquitin ligase complex, SLX5, together with DOA1, brings this aspect of protein degradation pathway into daily temporal regulation in yeast^64^. Although most of these functional classes are more or less expected, if one were using function as a predictor it would be impractical to sort through the hundreds of candidates with similar function based on e.g. GO categories. The PREMONition network suggests which proteins/molecules could be involved in the yeast clock.

In summary, we have developed a template for building interconnected networks that span multiple functional levels. The PREMONition algorithm depends on combining defined input sets of function-specific experimental results and non-specific, evidence-based, curated database entries. By incorporating the totality of interaction data, not just genetic interactions, we can move beyond transcription to different functional levels, achieving a molecular network that reflects the complexity of biological mechanisms.

## Materials and Methods

### Network construction

The network construction method consists of reading in pairs of molecules (as referenced by protein/gene names) and their interaction (ppi) probabilities that meet a set threshold to generate a network. This is done in a recursive manner, starting with a single protein, a seed, which grows into a network until no additional connections can be made. Then, the next protein is seeded etc. During this process the method is able to integrate NRCs in order to reach out to molecules encoded by rhythmic genes.

For this study, only one NRC between two rhythmic genes/proteins was allowed hence a last step was added to remove excess connections. After network construction, some proteins are highly connected. This makes the visual investigation of specific pathways difficult. We thus developed a routine to reduce complexity whilst maintaining the overall integrity of the network. Hence, for each connection, we asked “are all proteins in the network still connected if this connection is removed?”. If all proteins remain connected, then the connection is redundant and not required for network integrity^74^. If not, the connection is maintained. The result is a network with a minimal number of connections. The remaining connections have the highest interaction probability score. This process is used for visualization only and the connectivity of a protein was determined using the unprocessed network.

Known protein-protein and inferred functional interactions were obtained from the STRING interaction database (“protein.links.detailed.v9.1.txt”^75^). Each interaction is associated with an interaction probability score as derived from different information sources. To create a molecular interaction network, all interactions for a given organism that met a given interaction threshold were extracted. Taxonomy numbers used were; Hs:9606, Dm:7227, Nc:5141, and Sc:4932.

The ccgs were used for network construction based on the interaction confidence scores provided by STRING. In this study, an interaction confidence score with a threshold of 0.6 was assigned for *Neurospora* and 0.7 for human, *Drosophila* and yeast networks.

### Network analysis

In order to analyse the (re)constructed molecular interaction networks for functionality we assessed the enrichment of Gene Ontology (GO) annotations within the network using a hypergeometric test, with all p-values adjusted using the Benjamini and Hochberg method to correct for multiple testing. This is comparable to the GOrilla method^76^. A False Discovery Rate of 0.05 was maintained.

The most complete GO annotations for any organism typically reside in organism-specific databases supported by the community. Hence GO annotations were obtained from the Broad Institute for *N. crassa (https://www.broadinstitute.org*), Flybase for *D. melanogaster (http://flybase.org),* GOC for human (*http://www.geneontology.org*), and the *S. cerevisiae* Database (*https://www.yeastgenome.org*).

For the 1,000 yeast Monte Carlo GO experiments, the YeastMine API was used^77^. All ORFs, both verified and uncharacterized, were included in the background. Holm-Bonferroni multiple-testing correction^78^ was applied. All genes with a corrected p-value <= 0.05 were considered significant.

### Homology analysis

In order to use the InParanoid8 database, several ID-mapping steps were performed. Two hundred and sixty-nine *N. crassa* STRING identifiers were converted to NCBI Gene IDs (using NCBI genome.gff). These Gene IDs were subsequently mapped to Uniprot identifiers using the Uniprot mapping tool. For each organism (human, N.c and D.m) STRING identifiers had to be converted to Uniprot identifiers. For N.c, 269 STRING IDs were translated to NCBI Gene IDs; these were further converted by applying the Uniprot mapping tool to 283 yield unique refSeq protein IDs. 246 unique D.m. STRING IDs were converted to 219 refSeq protein ID (using Flybase) which then were converted to 267 unique Uniprot protein IDs. Finally, 258 unique H.s. STRING ID’s were translated into 245 unique Uniprot protein ID’s.

All orthologous protein pairs, including orthologous clusters identified by Inparanoid 8 (information downloaded on 25-March-2018), *i.e*. no actual filter, was used to identify orthologs between all three species. Enrichment of orthologous molecules within the networks was assessed using Phyper^79^ (Package stats version 3.2.2), a method to perform the hypergeometric test, from which a p-value <0.05 indicates significant enrichment.

### Network construction

The network reconstruction tool and GO analysis method were both written in the programming language Perl (version 5.8.8). Reconstructed network visualizations were made in Cytoscape (version 3.2.1)^80^.

### Yeast Culture Conditions

To reveal genes and processes involved in the regulation of the circadian clock of *S. cerevisiae*, a microarray experiment was performed according to methods described earlier^7^. Samples were taken from fermenter cultures (1 × 10^9^ yeast cells) every 3 hours over the first day of constant conditions (free-run), starting 1h before the temperature transition from 21°C to 28°C. Cells were centrifuged for 1 minute at 4°C and 14000 rpm. Pellets were flash-frozen in liquid nitrogen and stored at −80°C until processing.

### RNA Preparation

Total RNA was obtained using a modified version of the hot phenol method^81^. The flash-frozen yeast pellets were suspended in 400 μL AE buffer (50 mM NaOAc pH 5.3 and 10 mM EDTA) to which 40 μL SDS (10%) and 400 μL acidic phenol were added. Next, cells were disrupted by vortexing, followed by heating the sample to 65 °C for 30 min. The disrupted cells were quickly chilled on ice, centrifuged, and the aqueous phase was re-extracted using 400 μL acidic phenol followed by the addition of 400 μL chloroform, mixing, centrifuging and re-extraction of the liquid phase. The final step was purification and concentration of the RNA samples using the NucleoSpin RNA II kit (Macherey-Nagel). The RNA was used for transcriptome analysis (Service XS Leiden; Affymetrix yeast genome 2.0 arrays). Data can be accessed via Gene Expression Omnibus ID: GSE122152.

### Gene expression

Microarray data was analysed using the functional programming language R (version 2.15) and Robust Multi-array Average (RMA) normalization^82^ (Bioconduction “affy” package, version 1.34.0). The normalized expression values were analysed for rhythmicity with a 24h period using CircWave^83^ (http://www.euclock.org/results/item/circ-wave.html). This method performs an F-test on the fitted harmonics and calculates a *p*-value. We used only highly significant genes with a *p*-value less than 0.01. The thresholds used for the selection of rhythmic genes (‘ccgs’) in human, fly, fungus and yeast are listed in Table S6.

### Light sensitivity assay

The knockout library was obtained from EUROSCARF^84^ (Frankfurt am Main, Germany). Selected knockout strains were individually cultured overnight in 150 μl YPD (1% yeast extract, 2% peptone, 2% dextrose) medium in one well of a 96 well plate. Plates were held at 30°C with shaking at 160 RPM. The following day, the cultures were placed at 4°C until further use.

On the day before initiating the growth assay, 10 μl from each well of the 96-well plates were inoculated into 6 ml of YPD medium. The cultures were grown overnight at 30°C with shaking at 160 RPM. Prior to making the final dilution series, the cell number in the overnight cultures was adjusted such that the OD_600_ was 0.1. From this cell suspension, a series of five 1:3 dilutions were made in YPD. From each dilution, 2 μl was spotted on two independent YPD plates containing 2% agar. The knockout strains were compared to their parental wild-type strain (BY4742). Of the two, replicate plates containing replicate sets of dilution series, one was wrapped in aluminum foil (dark control) and then both were placed in the experimental setup. The dark controls thus control for small temperature effects. The light source consisted of 500 lux blue light LEDs (Bartheleme product # 1484330, 12V DC). Incubations were carried out in an incubator with the temperature (measured at the level of the plates) at 15.0 °C. Plates were incubated for four days, after which their images were saved (BioRad, ChemiDoc MP Imaging System) for analysis.

The scans of the plates were analyzed using an in-lab developed Python routine that determines the amount of growth *i.e.* the pixel intensity and density of each spot within a circle of a fixed radius. All pixels within the circle having a pixel intensity higher than a given background BRG-value are counted and used as a quantitative measure for colony formation. Data were analysed by the following method. The dilution where 25% growth of BY4742 control strain was determined for both light and dark incubation conditions (the value was interpolated from the growth-curves). For each strain, the LOG(1/(Light / Dark ratio)) was calculated. This conversion leads to an intuitive (lower) value for growth repression relative to control values.

## Acknowledgements

We thank members of the Merrow lab, Marco Grzegorczyk and John Hogenesch for comments on the manuscript. We thank Margriet van der Pol and Edwin Eelderink for technical assistance. This work was supported by the NWO (Open Programma, 820.02.014), the European Commission (FP6 EUCLOCK), the Rosalind Franklin Fellowships of the University of Groningen and LMU Munich.

## Author Contributions

J.B. and M.M. conceived the project, analysed data and wrote the paper. J.B. and Z.C.-E. performed experiments. E. L. analysed data and contributed to writing the paper.

